# Transcriptional shift and metabolic adaptations during *Leishmania* quiescence using stationary-phase and drug pressure as models

**DOI:** 10.1101/2021.12.03.470940

**Authors:** Marlene Jara, Michael Barrett, Ilse Maes, Clement Regnault, Hideo Imamura, Malgorzata Anna Domagalska, Jean-Claude Dujardin

## Abstract

Microorganisms can adopt a quiescent physiological condition which acts as a survival strategy under unfavourable conditions. Quiescent cells are characterized by slow or non-proliferation and deep down-regulation of processes related to biosynthesis. Although quiescence has been described mostly in bacteria, this survival skill is widespread, including in eukaryotic microorganisms. In *Leishmania*, a digenetic parasitic protozoan that causes a major infectious disease, quiescence has been demonstrated, but molecular and metabolic features enabling its maintenance are unknown. Here we quantified the transcriptome and metabolome of *Leishmania* promastigotes and amastigotes where quiescence was induced *in vitro* either through drug pressure or by stationary phase. Quiescent cells have a global and coordinated reduction in overall transcription, with levels dropping to as low as 0.4% of those in proliferating cells. However, a subset of transcripts did not follow this trend and were relatively upregulated in quiescent populations, including those encoding membrane components such as amastins and GP63 or processes like autophagy. The metabolome followed a similar trend of overall downregulation albeit to a lesser magnitude than the transcriptome. Noteworthy, among the commonly upregulated metabolites were those involved in carbon sources as an alternative to glucose. This first integrated two omics layers affords novel insights into cell regulation and shows commonly modulated features across stimuli and stages.

## 1. Introduction

*Leishmania spp*. are parasitic protozoa with two overarching life-cycle stages. Extracellular flagellated promastigotes reside in the gut of the sandfly vector, where the proliferative procyclics differentiate into the infective but non-proliferative metacyclics that are transmitted during the sandfly bite [1]. Within the mammalian host, the metacyclics enter macrophages where they differentiate into ovoid aflagellated amastigotes adapted to live in the parasitophorous vacuole of phagocytic cells. Within this inhospitable environment, *Leishmania* amastigotes activate mechanisms that allow them to survive and proliferate and propagate the infection [2]. However, in addition to entering a slowly proliferating phase characterised by a stringent metabolism [3], subpopulations can enter a non-proliferative quiescent state [4–6]. Understanding of quiescence in *Leishmania* is of great significance given the roles the process probably plays in chronic and asymptomatic infections and the recrudescence of the disease after therapy without associated development of drug resistance [7–10].

Quiescence is a reversible condition in which a cell does not divide but retains the potential to re-enter the replicative state [11]. Quiescent cells emerge spontaneously or at enhanced rates in response to different stresses such as drug pressure, nutrient limitations, or host immunity [12–16]. During quiescence, cells modulate their molecular processes and metabolic pathways to save energy and shut-down metabolic pathways that may be vulnerable to stress responses or provide products that can stimulate effective host immunity. [12, 17]. Biosynthesis of macromolecules is reduced but processes important for survival such as carbon storage, autophagy, and host manipulation are activated [12, 17]. In *Leishmania*, quiescence has been reported in three species: *L. mexicana*, *L. major*, and *L. braziliensis* [4–6]. It has also been inferred in *L. donovani* [18]. In *L. mexicana* and *L. braziliensis* quiescent amastigotes showed downregulated synthesis of ATP, ribosomal components and proteins and alterations in membrane lipids [5, 6, 19]. These preliminary reports allowed the development of negative biomarkers such as the expression of GFP integrated into the ribosomal locus, which allows active cells to be distinguished from quiescent ones *in vitro* and *ex vivo* [6]. However, in *Leishmania*, several key knowledge gaps remain, including understanding whether quiescence can be induced by drug pressure, how quiescence is maintained, and the degree to which quiescent cells derived through different means may share molecular and metabolic traits in response to different environmental insults.

Here we undertook a first integrated transcriptomic and metabolomic study of quiescence in two life stages of *L. lainsoni* (promastigotes and amastigotes), upon *in vitro* induction by stationary phase or drug pressure. The integrations of two quiescence models and two life stages allowed us to identify modulations independent of the environmental insults and the parasite stage. These core changes in quiescent *Leishmania* parasites may represent an essential set of traits that allow their survival. Moreover, these core changes could also point to the molecular and metabolic landscape appearing in other forms of quiescence and provide positive markers to track quiescence and targets to enable pharmacological disruption of the process.

## 2. Results

### 2.1. PAT drug pressure and stationary phase as models of quiescence

In several microorganisms, a quiescent state can be induced by stressful conditions such as drug pressure or changes associated with the stationary phase of the growth curve, such as nutrient limitation and accumulation of waste products, including acidification of the medium [13, 15]. Here we implemented these two distinct models in which parasites become quiescent for the further molecular characterization of quiescence. The advantage of integrating molecular and metabolic traits in two quiescence models is that it discriminates changes inherent to a quiescent condition from changes related to the stimuli. We chose Potassium antimonyl tartrate (PAT) because: i) it represents the active form of a drug still used as first-line treatment in many endemic areas, and ii) there is striking evidence of therapeutic failure without the development of drug resistance [10, 20]. *L. (Viannia) lainsoni* clone PER091 EGFP Cl1 was used as a model because it has been adapted to grow axenically as amastigotes a feature that may be challenging for *L. (V.)* species. To monitor proliferation, growth curves in the absence or presence of PAT were monitored microscopically. Quiescence in viable cells was tracked by quantifying the rEGFP expression: its expression is high in proliferative and metabolically active cells and substantially downregulated in non-proliferating cells with stringent metabolism [6].

The growth curves showed that in the absence of PAT both promastigotes and amastigotes had an exponential increment in cell density during the first two days of growth *in vitro*, reaching a stationary phase by day 4 (Figure 1a). In both stages, a continuous PAT exposure (1 μg/mL, ~10 fold IC50) had a negative effect in their proliferation, causing a substantially decreased or arrested growth from day 1, with this repression maintained over the following 5 days (two way ANOVA, Stage; *p* = 0.15, PAT *p* = 2.8 × 10^-6^). Besides an apparent cytostatic effect, PAT was also cytotoxic, and cell viability was ~77.2 and ~47% after 2 days of exposure to 1 μg/mL of PAT in promastigotes and amastigotes respectively. After enrichment for viable cells using density gradients, cell viability rose to ~87 and ~90 % for promastigotes and amastigotes respectively (despite several attempts, higher enrichments were not feasible). When cells were placed in a fresh medium without PAT pressure, both *Leishmania* stages resumed their proliferative condition. The reversibility was also tested with *Leishmania* cells recovered from 9 μg /mL of PAT pressure (~90 fold IC_50_), indicating that at least a fraction of the whole population endures pharmacologically relevant concentrations of PAT. The regrown proliferative population emerging after the release of the 9 μg /mL of PAT exposure retained high susceptibility to PAT (IC50 ~ 0.05 μg / mL), similar to the population without exposure to drug pressure ruling out that the survival was due to a selected drug resistance mechanism

**Figure 1.**
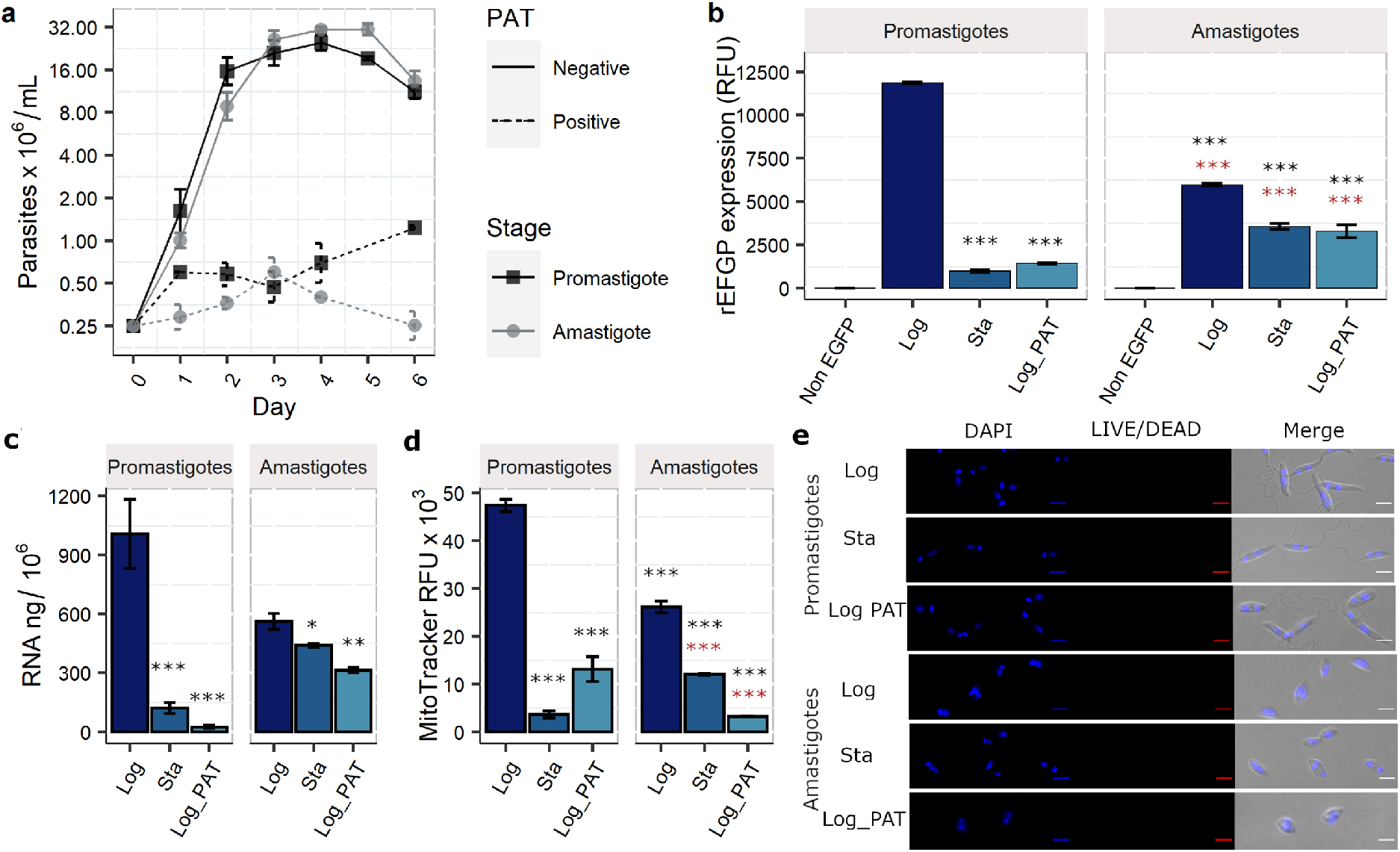
Evaluation of stationary phase and drug pressure as models of quiescence. Four parameters with associated downregulation in quiescent cells in addition to cell viability were evaluated. (a) Assessment of proliferation as indicated by growth curve of promastigotes and amastigotes in absence (solid lines) and presence (dashed lines) of Potassium Antimonyl tartrate (PAT). (b) rEGFP expression in proliferative (Log) and non-proliferative (Sta, Log_PAT) conditions. (c) Quantification of the total RNA content per million cells. (d) Quantification of the mitochondrial membrane potential as indicated by the fluorescence of Mitotracker. In figures b-d, the results represent the mean ± SEM of three biological replicates. The asterisks represent statistically significant differences after Tukey post hoc test; *p <0.05, **p <0.01, **p <0.001. The asterisks in black and red represent statistically significant differences compared to promastigotes and amastigotes in Log respectively. e, Confocal images of Leishmania cells under the evaluated conditions (after enrichment of viable cells for condition Sta and Log_PAT). The pictures show their morphology, size, and cell viability as indicated by their membrane integrity and absence of cytoplasmatic staining with the non-permeable dye LIVE/DEAD^™^ Fixable Red Stain. The maintenance of their nuclear and kinetoplast DNA is shown with DAPI staining. The sale bar represents 5 μM.

For each life stage, the expression of rEGFP was monitored by flow cytometry in a replicative condition represented by parasites harvested in the exponential phase of the growth curve (Log, no drug pressure) and two non-proliferative conditions: parasites harvested in the stationary phase (Sta, no drug pressure) and parasites in the exponential phase of the growth curve that have arrested their growth in response to PAT drug pressure over 48 hrs (Log_PAT) (Figure 1b). In promastigotes the rEGFP expression dropped from 11.8 × 10^3^ relative fluorescent units (RFUs) in Log to 1.0 × 10^3^ (~ 8 %) and 1.4 × 10^3^ (~12%) in Sta and Log_PAT, respectively. In amastigotes the rEGFP expression decreased from 5.9 × 10^3^ in Log to 3.6 × 10^3^ (61%) and 3.3 × 10^3^ (56 %) in Sta and Log_PAT, respectively (two-way ANOVA, Stage *p*= 1.0 × 10^-14^, Condition *p* = 8.3 × 10^-08^). The arrested growth and the decreased levels of rEGFP expression indicated that both *Leishmania* stages in conditions Sta and Log_PAT had adopted a quiescent state.

### 2.2. Evaluation of additional quiescence indicators: total RNA content and mitochondrial activity

Quiescent cells downregulate their total RNA content and reduce mitochondrial activity [21, 22]. We also analysed these two parameters to monitor quiescence in conditions Sta and Log_PAT post enrichment of viable cells. The total content of RNA dropped dramatically in both conditions, as indicated by the fluorometric quantification of the isolated RNA (Figure 1c). In promastigotes, the RNA content/ 10^6^ cells dropped significantly from ~1010 ng in Log to 121 ng (~12 %) and 24 ng (~2.4 %) in Sta and Log_PAT, respectively.In amastigotes, the RNA content/ 10^6^ cells decreased from 563 ng in Log to 440 (78.6 %) and 314 (55.0 %) in Sta and Log_PAT respectively (two-way ANOVA, Stage *p*= 2.1 × 10^-6^, Condition *p* = 6.9 × 10^-8^). In most eukaryotes, rRNAs are the most abundant component of total RNA (up to 80 %). Gel electrophoresis of total RNA indicated that this is also the case in *Leishmania*, and the rRNAs remained the most abundant RNAs in Sta and Log_PAT cells (Figure S1).

Mitochondrial activity was evaluated by the MitoTracker RFU (Figure 1d). In promastigotes the signal dropped from 47.3 ×10^3^ RFU in Log to 3.6 × 10^3^ (7.6 %) and 13.1 × 10^3^ (34.4 %) in Sta and Log_PAT respectively. In amastigotes the MitoTracker RFUs decreased from 26.1 ×10^3^ in Log to 12.4 × 10^3^ (47.5 %) and 3.2 × 10^3^ (12.1 %) in Sta and Log_PAT respectively (two-way ANOVA, Stage *p* = < 0.0001, Condition *p* < 0.0001). Under the evaluated quiescent conditions, viable cells did not demonstrate substantial changes in their body size compared to the proliferative condition (Figure 1e). The following experiments outline the transcriptomic and metabolic characterization of both quiescent conditions compared to proliferative cells of each *Leishmania* life-cycle stage as non-quiescent controls. An outline of the experimental setup and analysis is in Figure S2.

### 2.3. Evaluation of the mRNAs transcriptome size and global shift in the levels of mRNAs

The sum of all mRNA molecules per cell in this article will be referred to as transcriptome size. While a global shift in the mRNA transcriptome refers to a change in the average number of individual mRNA molecules per cell, which follows a similar trend across most genomic locations in the whole genome. In *Schizosaccharomyces pombe*, quiescent cells diminish their transcriptome size for RNA polymerase I transcribed rRNAs and RNA polymerase II transcribed mRNAs, to 11 and 18 % the levels in replicative cells, respectively [23]. Similarly, *Leishmania* quiescent amastigotes diminish their expression and content of rRNAs [19], a finding also reported here in response to both stationary phase and PAT pressure. In order to evaluate changes in the transcriptome size for mRNAs, we performed bulk RNAseq coupled with ERCC spikes-in (these are a set of external non-*Leishmania* RNAs that can be used to calculate a per-sample normalization factor (NF), which corrects the inherent variation of the total RNA content/cell under different conditions and the errors accumulated during the library preparation and sequencing [24], Figure S3 a, b). This approach allows robust identification of changes in transcriptome size and the identification of transcripts with altered abundance (number of mRNAs per cell) when comparing different conditions [25]. The samples had a good correlation between the expected and experimental fold changes for the ERCC spikes-in, which is indicative of their good performance in our experimental setup (Figures. S3 c, d). After normalization, principal component analysis (PCA) showed that most of the variation in the number of mRNAs per cell (PC1, 98%, PC2 1%) was explained by the experimental condition: parasites in Log_PAT are most dissimilar to parasites in Log (for both promastigotes and amastigotes (Figure 2a)). The stage or condition of the parasite did not have an effect in the number of genes being expressed with an average of 8,014 out of the 8,295 annotated CDS in the reference genome being detected (two-way ANOVA, Stage *p* = 0.96, Condition *p* = 0.15). Of those genes not detected 34% were annotated to encode putative proteins. The results also indicated that the mRNA transcriptome size undergoes a dramatic decrease in quiescent cells compared to proliferative cells (Figure 2b). In promastigotes the sum of normalized mRNAs dropped from 1.8 × 10^8^ in Log to 2.1 × 10^7^ (~11.7 %) and 7.2 × 10^5^ (~0.4 %) in Sta and Log_PAT respectively. In amastigotes the sum of normalized mRNAs decreased from 1.4 × 10^8^ in Log to 6.1 × 10^7^ (~45.1 %) and 7.9 × 10^6^ (~5.8 %) in Sta and Log_PAT respectively (two-way ANOVA, Stage *p* = 0.96, Condition *p* < 0.0001). We then evaluated the impact of diminished mRNA transcriptome size in the levels of each mRNA encoded in the genome by comparing quiescent cells against proliferating cells. On each dichotomic comparison, the vast majority of mRNAs had decreased levels indicating the occurrence of a global down-shift in the mRNA transcriptome of quiescent cells (Figure 2c, Table S1). The abundance of 1,509 transcripts (18.2 % of all transcripts) was decreased in all quiescent cells across conditions and stages when compared to proliferative cells (cutoff log2 FC < −1 and BH adjusted *p* value <0.05). The number of decreased transcripts was 7,847 (94.6 %) if only the overlap between quiescent promastigotes (Sta and Log_PAT) and amastigotes under PAT pressure were considered (Figure 2d). Quiescent cells across stages and conditions had a negligible number of mRNAs with increased abundance compared to proliferative cells, and none of them were shared across conditions (Figure 2d). We also evaluated if the impact of the transcriptome down-shift was uniform or differential at the level of individual mRNAs by calculating the correlation between the mRNA transcriptome of proliferative and quiescent cells. A very similar effect over most mRNAs in the transcriptome was indicated by the high pairwise correlations between quiescent vs. proliferative cells (r values ranged from 0.86 in Pro Log_PAT to 0.97 in Pro Sta, Figure 2 c-d). The results here show that during quiescence, *Leishmania* has a diminished transcriptome size and global down-shift.

**Figure 2.**
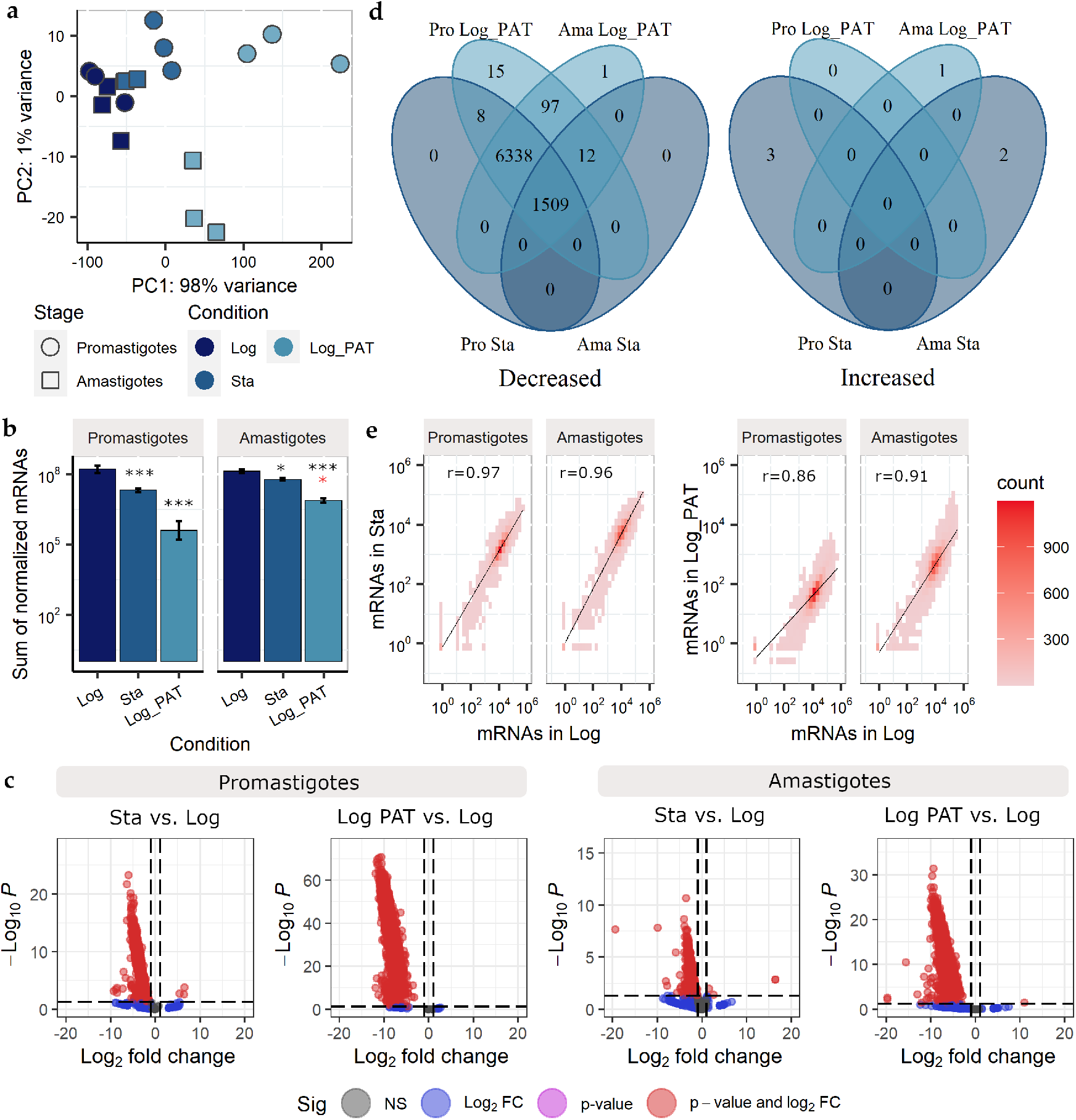
mRNA transcriptome size and global down-shift during quiescence. a, Principal component analysis showing that most of the variability in the abundance of the mRNAs can be explained by the proliferative condition of Leishmania with quiescent cells induced by PAT drug pressure being the most dissimilar compared to proliferative cells in both promastigotes and amastigotes. b, Sum of normalized mRNAs in each condition and stage of Leishmania showing the diminished transcriptome size in quiescent cells compared to proliferative cells. The bars represent the mean ± SE of three biological replicates. The asterisks in black color represent statistically significant differences after the Tukey post hoc test; * indicates p <0.05, indicates *** p <0.01. Black and red colors represent statistically significant differences in quiescent cells compared to promastigotes and amastigotes in Log, respectively. c, Volcano plots showing the magnitude of change in the relative abundance of each mRNA in quiescent cells compared to proliferative cells. The plots show the global downshift of the transcriptome in quiescent cells induced by both PAT or stationary phase in both Leishmania stages as most of the mRNAs have substantially decreased levels and are located in the top left side of the plots. On each plot, the difference is represented by the log2 fold change between two conditions, and its significance is represented by the negative logarithm of the p value. The two vertical lines in each plot represent the log2 fold change cutoff of 1 and −1 for increased or decreased levels, respectively. The horizontal line represents a 0.05 cutoff for the p value. d, Venn diagram representing the number of mRNAs with decreased or increased abundance in quiescent cells compared to proliferative cells. e, Pearson correlation between the abundance of mRNAs in quiescent conditions compared to proliferating cells. The color indicates the number of mRNAs within a particular bin.

### 2.4. Evaluation of the transcriptome composition

Although much lower, the absolute abundance of most of the mRNAs in non-proliferative cells was generally highly correlated with the abundance of mRNAs in proliferative cells. However, the abundance of some specific mRNAs did not follow the global trend, indicating that the proportion of individual mRNAs relative to their transcriptome (which we will refer to as transcriptome composition) remained unchanged for most of the mRNAs but with changes for those not following the global trend. To evaluate the transcriptome composition, the transcripts per million of reads (TPM) of each mRNA in all conditions and each stage of the parasite were calculated. When using TPM, the sum of all TPM values in each sample is the same. This makes it easier to calculate the proportion of each mRNA in relationships to its own transcriptome and allows further comparison between samples [26]. The PCA showed that most of the variability in the transcriptome composition was explained by the biological condition of the parasite. For conditions Log and Sta, both promastigotes and amastigotes clustered very closely. For the Log_PAT condition, each stage may be considered as an independent cluster (Figure 3a). To identify the differentially modulated transcripts, we compared the TPM values of quiescent promastigotes and amastigotes against their respective proliferative condition (Table S2a). The number of transcripts with modulated TPMs ranged from 800 (~ 9.6 % of all transcripts) in stationary promastigotes to 2,239 (~ 26.9 %) in amastigotes under PAT pressure. The number of transcripts and their directional modulation in each stage and condition are shown in Figure 3b. When compared to proliferative cells, both *Leishmania* stages under PAT drug pressure revealed a higher number of modulated transcripts (Pro=1,703, Ama= 2,239) than corresponding stationary cells do (Pro=800, Ama=1,488), with the largest fraction showing downregulated TPMs (Pro=1,383, Ama= 1,799). This indicates that both stages under PAT drug pressure have additional suppression for a set of transcripts beyond the overall transcriptome down-shift. The further data integration identified a total of 135 mRNAs (down=87, up=48) that were modulated and shared across quiescent conditions and stages of *Leishmania* (Figure 3.c).

**Figure 3.**
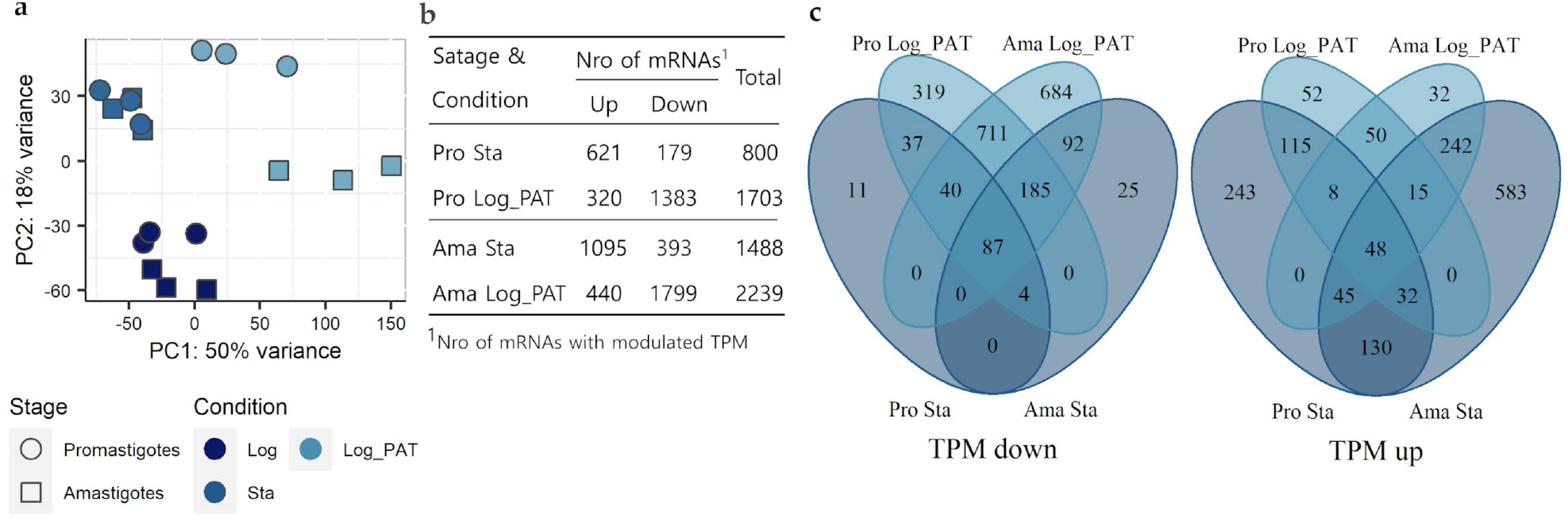
Transcriptome composition and its modulations during quiescence. a, Principal component analysis shows quiescent cells induced by stationary phase and PAT pressure differ in their transcriptome composition compared to proliferating cells in both promastigotes and amastigotes. b, Summary of mRNAs modulated in each Leishmania stage and condition. c, Venn Diagrams showing the number of mRNAS with modulated TPMs in the conditions Sta and Log_PAT compared to Log in each Leishmania stage. Pro, promastigotes; Ama, Amastigotes.

These transcripts are candidates for involvement in quiescence irrespective of *Leishmania* developmental stage or the stimulus triggering the phenotype (Figure S4). Additional information about their abundance relative to other transcripts (TPM distribution) and broad genomic location is shown in Figure S5. Besides the core set of mRNAs modulated in all quiescent conditions, the set of mRNAs modulated in both promastigotes and amastigotes but specific to the stationary phase (conditions STA: down=0, up=130) or the drug pressure (Log_PAT: down=711, up=50) are shown in (Table S2 b). These sets of modulated transcripts are likely essential and involved in the mechanisms that allow quiescent cells to adapt to the special needs encountered during these different stressful environments.

### 2.5. GO and GSEA analysis

To evaluate which biological features are altered during quiescence, we performed functional enrichment analysis, also known as overrepresentation analysis (ORA). The ORA and its network analysis for the core set of mRNAs modulated in all quiescent conditions and stages revealed several biological changes as indicated by many significantly enriched gene ontology (GO) terms corresponding to several cell processes/components (Table S2b, Figure 4). Among the downregulated biological processes in quiescent cells were mitochondrial transmembrane transport and ATP biosynthesis in the mitochondria, including the cellular components electron carrier cytochrome c and the complex cytochrome c reductase as well as several subunits of the ATP synthase complex, being among the most representant transcripts: ATP synthase F1, ATP synthase alpha, ATP synthase beta and ATP synthase gamma. Within the nucleus, the process of regulation of DNA replication and components of the DNA packaging complex were also downregulated in quiescent cells. Within the category DNA replication, the transcript encodes the proliferative cell nuclear antigen (PCNA), an essential positive regulator of nuclear DNA replication [27, 28]. For DNA packaging, the transcripts encoded copies of histones H2A and histones H2B. The translation process is inferred to be diminished since transcripts encoding three elongation factors: elongation factor 1-beta, elongation factor 1-gamma and elongation factor 2 were all downregulated. For the list of genes with upregulated TPM values, the processes of autophagy and amino acid catabolism via the Ehrlich pathway were enriched (Figure 4). Two copies of the gene ATG8/AUT7/APG8/PAZ2 encoding an ubiquitin-like protein that has a plethora of roles during the biogenesis of autophagosomes from early stages until the final event of fusion with lysosomes were upregulated [29]. In the process of catabolism of amino acids via the Ehrlich pathway the gene encoding a putative alpha keto acid decarboxylase was upregulated. This pathway is a source of alpha keto acids such as pyruvate during conditions of nutritional starvation [30]. The biological process, “modulation of process of other organism”, represented by genes encoding a major metalloendopeptidase called Leishmanolysin/GP63 was also upregulated. It should be highlighted that among the core set of upregulated transcripts, a high proportion had membrane localization (n=12, ~25%), including 4 copies for a “surface antigen-like protein”. Another considerable proportion of transcripts were encoding hypothetical proteins (n=19, ~40 %). Therefore, it is not yet possible to infer a biological function.

**Figure 4.**
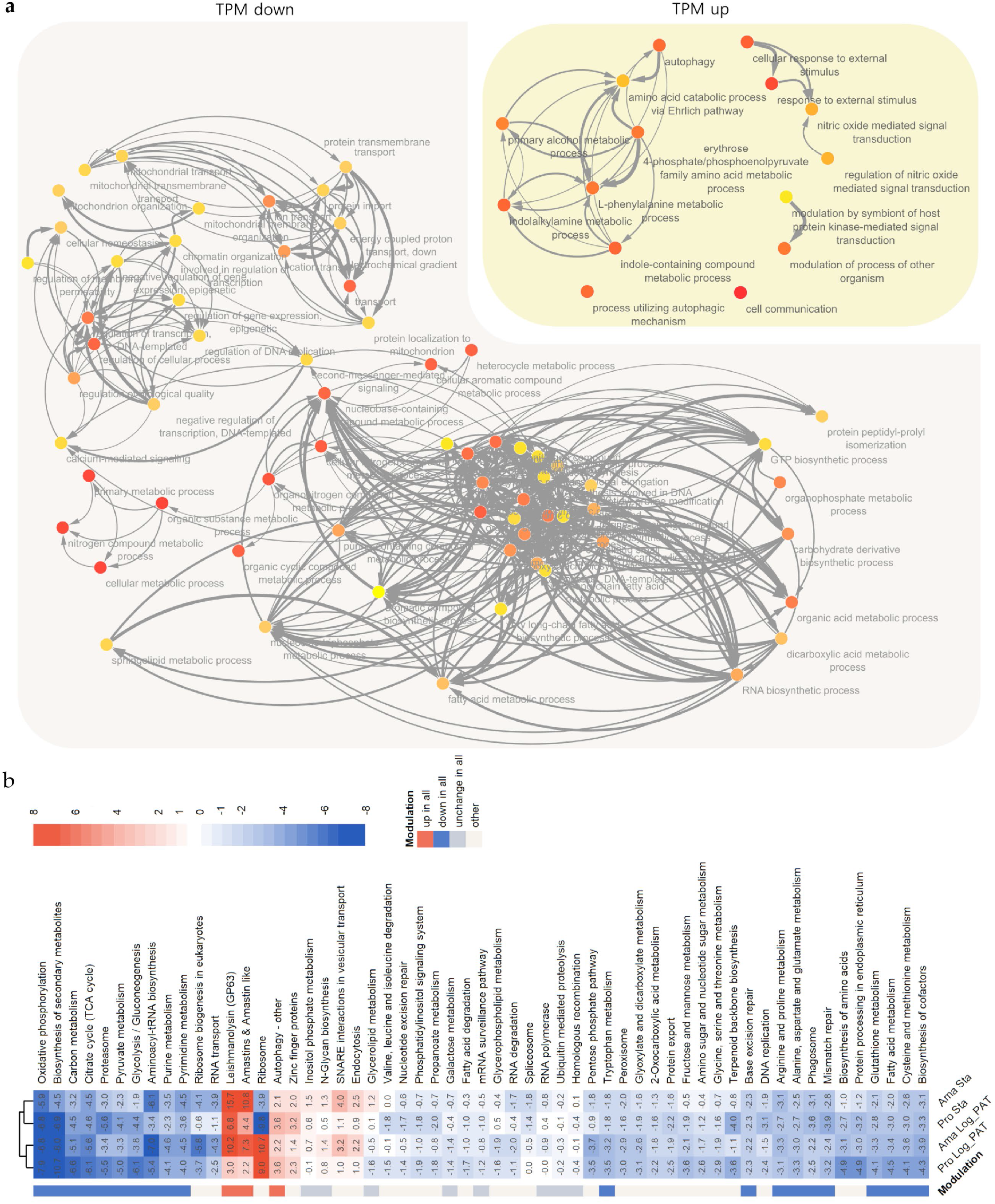
ORA and GSEA analysis. a, ORA and network analysis for the core set of mRNAs modulated and shared among all quiescent conditions. ORA indicates many biosynthetic processes are downregulated while few processes related to ‘cell host modulation’ and the recycling of nutrients are upregulated. The color of the bubble from yellow to orange indicates more significant adjusted p values (cutoff adj *p* <0.05). b, GSEA shows the major set of mRNAs upregulated and shared among all quiescent cells correspond to: leishmanolysin (GP63), amastin and amastin-like proteins, and autophagy-related genes. On the contrary, among other processes, the biosynthesis of several amino acids and biosynthesis of aminoacyl tRNA are downregulated in all quiescent cells regardless of the stimuli. The absolute value in each cell of the heatmap measures the magnitude of gene-set changes in log2 scale, and its sign indicates the direction of the change. The modulation of a particular set was considered up in all (red) or down in all (blue) if the p values were statistically significant in all conditions. While it was considered unchanged (grey) if the p value in all conditions were non-significant. Additional details are in Table S2 c-d

To complement the GO enrichment analysis, we also performed gene set enrichment analysis (GSEA), which does not analyse separate lists of upregulated or downregulated genes according to an pre-established cutoff, instead analysing all genes belonging to a functional category. GSEA showed similar results to ORA, indicating that many biosynthetic processes are significantly diminished in all quiescent conditions and stages and extended the results to the metabolism of amino acids such as alanine, aspartate, glutamate, arginine, proline, tryptophan, cysteine, and methionine. Among others, processes such as glycolysis/ gluconeogenesis, pyruvate metabolism, citric acid cycle (TCA cycle), fatty acid metabolism, and aminoacyl-tRNA biosynthesis were all down-regulated (Figure 4b, Table S2 d-e). Regarding the upregulated categories, GSEA confirmed that in addition to autophagy and *GP63* the expression of genes encoding amastins and amastin-like surface proteins were upregulated in all quiescent conditions (Figure 4b, Table S2d). In a stringent biological state of overall downregulation, unchanged pathways or processes across conditions may indicate they are essential for quiescent and proliferative cells. Among these were: inositol phosphate metabolism, N glycan biosynthesis, ubiquitin-mediated proteolysis, and homologous recombination.

### 2.5. Metabolomics

Metabolite identification and relative quantification was performed on the same cell samples as those used for transcriptomics, with an untargeted LC-MS approach. A total of 689 metabolites were putatively annotated. Quiescent cells had a diminished metabolome compared to proliferative cells, as indicated by the decrease in the sum of the total ion content (TIC) per equivalent amount of cells. In promastigotes the total signal decreased from 2.28 ×10^10^ in Log to 1.02 ×10^10^ (~44.8 %) and 0.99 ×10^10^ (43.6%) in Sta and Log_PAT respectively. In amastigotes the total signal decreased from 1.86 ×10^10^ to 0.63 ×10^10^ (~33.9 %) and 1.09 ×10^10^ (58.8 %) in Sta and Log_PAT respectively. The main factor for this decrease was the condition of the parasite (two-way ANOVA, Stage *p*= 0.068, Condition *p* <0.0001) (Fig 5. a). We then evaluated the metabolome with two approaches (i) changes in abundance of metabolites per equivalent number of cells (cell normalized data) and (ii) changes in the metabolome composition by normalizing the intensity of each metabolite to the TIC of its respective sample (IPT normalized data) (Table S3a, b). Both analyses gave complementary insights into metabolism during quiescence. The cell normalized data indicated quiescent cells have multiple metabolites of decreased abundance compared to proliferative cells (up to 166 in Ama Sta). In contrast, fewer metabolites had increased abundance (up to 44 in Ama Sta) (Figure 5b). However, when normalizing to the TIC, the metabolome composition revealed that quiescent cells have up to 56 metabolites (in Ama Sta) with decreased IPT levels and up to 277 metabolites (also in Ama Sta) with increased IPT levels (Figure 5b). A principal component analysis (PCA) showed that variability in metabolite abundance and composition were explained in large part by both the parasite stage separated by the PC1 (Cell 38 %, IPT, 42 %) and the parasite conditions separated by the PC2 (Cell 23%, IPT 22%) (Figure 5c). As indicated by the PCAs, many changes in metabolite levels and the metabolome composition in response to stressful stimuli were associated with the developmental stage of *Leishmania*. Therefore, the number of commonly modulated metabolites was higher between Sta and Log_PAT within the same stage than between Sta (or Log_PAT) in different *Leishmania* stages. However, a core set of metabolites were modulated across all quiescent conditions regardless of the stage (abundance: down=20, up= 8; composition: down=2, up=17; Figure 5d, e).Within the cell normalized dataset, quiescent parasites of both stages had lower abundance of several metabolites involved in the metabolism of amino acids (Figure 6). Nucleotide metabolism was also modulated with all detected nucleotides being decreased, while the abundance of some intermediary metabolites involved in the salvage or degradation of purines were increased. The abundance of several carbohydrates involved in the pentose phosphate pathway that leads to the synthesis of pentoses such as ribose 5-phosphate (precursor for the synthesis of nucleotides) were also decreased. While the abundance of carbon sources such as sucrose and 3 hexose-multimers (labelled as cellopentaose, cellohexaose and celloheptaose) were identified as increasing in quiescent cells. These penta, hexa and heptahexoses have previously been inferred [31] to derive from mannogen, a poly-mannose carbohydrate reserve used by *Leishmania* [32]. Interestingly, while most structural phospholipids were not modulated, a number of free fatty acyls and also sphingolipids were increased in quiescent promastigotes, while no clear trend was observed in amastigotes. When evaluating the composition of each metabolome (IPT normalized data), the trend observed with the cell normalized data set was maintained for nucleotide and carbohydrate metabolism. For quiescent cells, the metabolites within these categories had overall decreased IPT levels compared to proliferative cells. Additional information was also captured. For instance, uric acid that is the end product of purine degradation had increased IPT levels in quiescent amastigotes indicating they may have an enhanced purine nucleotides degradation (Figure S6). Within carbohydrates, potential carbon sources including mannose derivatives such as Mannose 6-phosphate and GDP-mannose had increased IPT levels in all quiescent cells compared to proliferative cells. Finally, all quiescent conditions showed that overall fatty acids had increased IPT levels (Figure S6).

**Figure 5.**
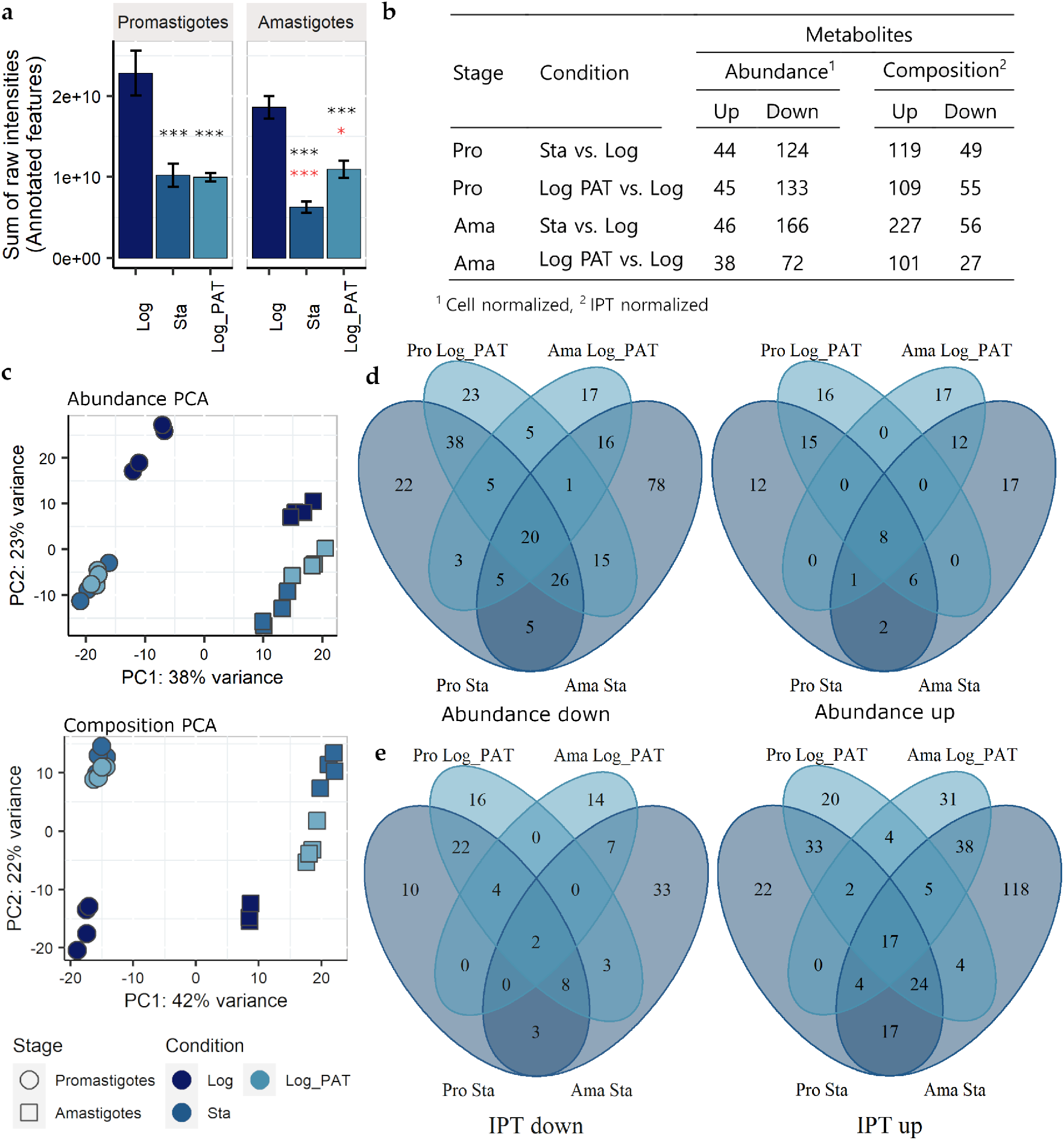
Quiescent cells have a distinct metabolic profile when compared to proliferating cells. The figures represent 689 putatively annotated metabolites. a) In all quiescent cells of each Leishmania stage, there is an overall downregulation in the levels of metabolites (either in condition Sta or Log_PAT) as indicated by the decrease in the sum of the peaks intensities of all annotated features constituting their metabolome. The asterisks in the top of each bar in black color represent significant differences after Tukey’s test; * *p* <0.05, ** *p* <0.001. The asterisks in black and red represent differences in comparison to promastigotes and amastigotes in Log respectively. b) Summary of the number of modulated metabolites in quiescent cells induced by condition Sta or Log_PAT in both promastigotes and amastigotes. The results after normaliz-ing by equivalent amount of cells or by IPT are shown (cutoff |log2 FC| >1 and BH adjusted *p* <0.05). c) Quiescent cells in both stages have a distinct metabolic profile as indicated by the clear separation of quiescent and non quiescent cells by the PCA 2. d-e) Venn Diagrams showing the relationship between the set of modulated metabolites (panel b) across the different condition and stages of the Leishmania. Pro, promastigotes; Ama, amastigotes.

**Figure 6.**
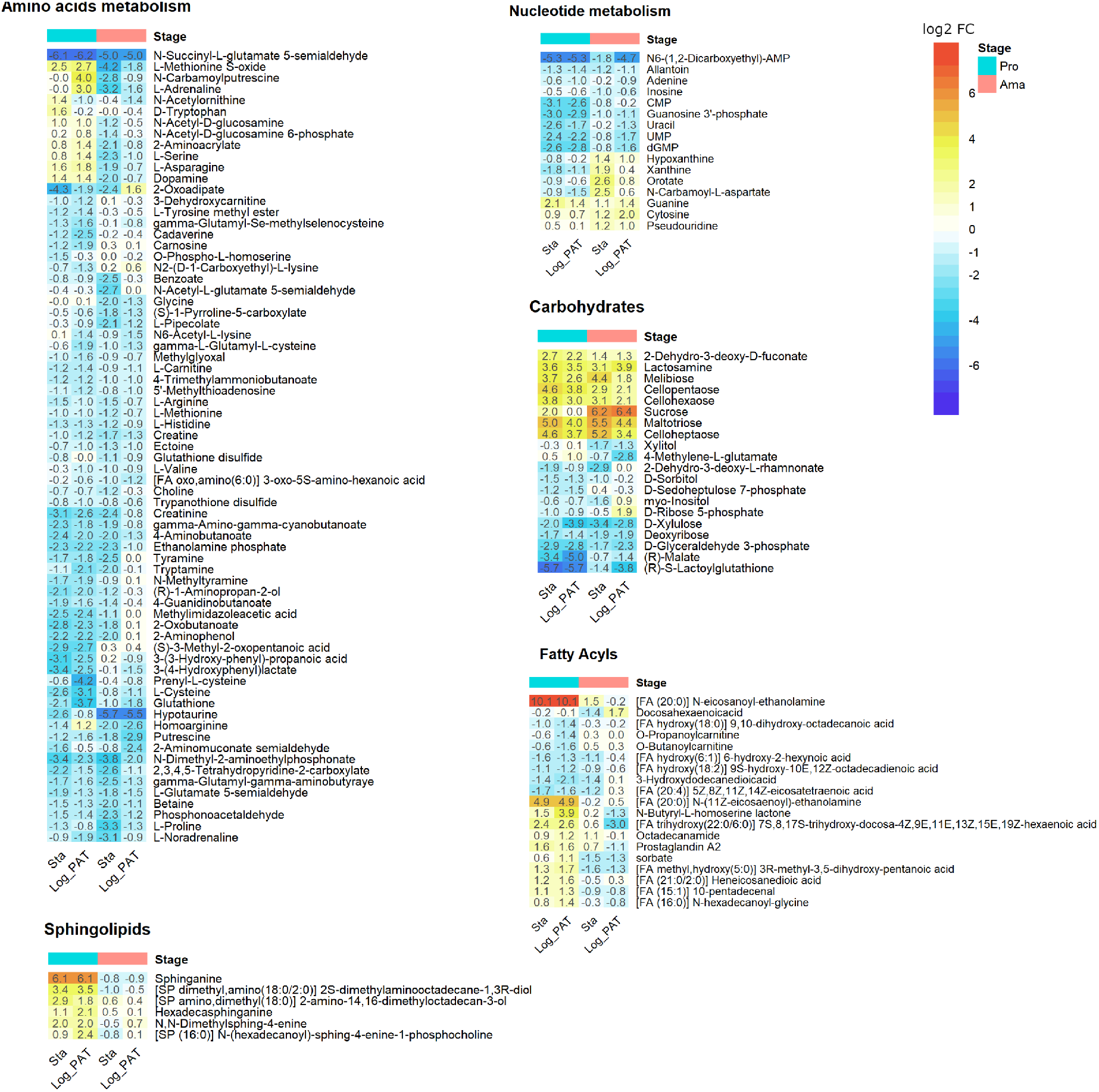
Main metabolic categories having metabolites with modulated abundance in quiescent cells induced by station-ary or drug pressure compared to proliferating cell. The numbers represent the log2 FC in comparison to condition Log (cell normalized dataset). Only metabolites for which at least one of the conditions has an |log2 FC| > 1 and a BH adjusted *p* <0.05 are shown. Pro, promastigotes; Ama, amastigotes.

## 3. Discussion

Quiescence in microorganisms describes a state in which cells alter their metabolic status and cease to divide. Quiescence appears to be an ancient evolutionary trait that enables cells to transiently lose susceptibility to environmental stresses that are harmful to proliferating cells; nowadays this includes insusceptibility to therapeutic drugs. In this study, we show that *L. lainsoni* can adopt a quiescent state in response to PAT drug pressure as well as when reaching the stationary phase in cell culture. We went on to characterise the transcriptomes and metabolomes of cells in both quiescent states and compared them with the same data-types derived from proliferative cells, using both the promastigote and amastigote stages of the parasite. We show *L. lainsoni* quiescent cells have multiple differences at both transcriptomic and metabolomic levels compared to their proliferative counterparts. Moreover, we found that quiescent cells exhibit common traits that are independent of the stressful stimuli associated with their derivation, namely: a diminished transcriptome size with global down-shift, considerable compositional changes in their mRNAs, and metabolic changes including a possible shift towards increased utilisation of the carbohydrate storage molecule mannogen.

At the transcriptomic level, we evaluated, for the first time in *Leishmania*, the differences in the transcriptome size in response to different conditions. The diminished transcriptome size with global down-shift during quiescence under PAT to 0.4 % and 5.8% of their levels of proliferative cells for promastigotes and amastigotes respectively suggest a general mechanism of transcriptional repression. A similar decrease in transcriptome size has been described in early quiescent cells of *Schizosaccharomyces pombe* after just 12 hrs of nitrogen deprivation (down to 18 % compared to proliferative cells) [23]. In *Saccharomyces cerevisiae*, fully differentiated quiescent cells revealed a massive 30-fold drop in their mRNA levels [33, 34]. The global transcriptional down-shift we note in *L. lainsoni* is also comparable to the decreased turnover of RNA reported during quiescence in *L. mexicana* amastigotes *[5]*. Our results suggest a central control mechanism regulating the overall transcription by RNA pol II at the level of initiation. We propose that during *Leishmania* quiescence a factor (or factors) are positively modulated in response to stressful stimuli to limit overall gene transcription. In *Saccharomyces cerevisiae*, histone deacetylation appears to play such a role during quiescence by rendering regions of transcription initiation less accessible to RNA pol II [34]. Of note is the fact that in *Leishmania*, like other Kinetoplastid protozoa, there is a widely held view that regulation of Pol II transcribed genes occurs at a post-transcriptional level with differential stability of mRNA responsible for much of the altered RNA levels throughout the life cycle. However, these cells do have histones and a dynamic process of histone post-translational modification. For instance, *L. donovani* predominantly shows euchromatin at transcription start regions in fast-growing promastigotes, but mostly heterochromatin in the slowly proliferating amastigotes, reflecting a previously shown increase of histone synthesis in the latter stage [35]. Another mechanism behind diminished RNA levels involves an alteration of the transcription machinery. For instance, in quiescent myoblasts, RNA pol II has a cytoplasmatic rather than nuclear localization [36]. Roles for these processes in the diminished transcriptome size associated with quiescence in *Leishmania* will be a topic of fruitful future research. A mechanism for transcriptional repression in quiescent cells is an inter-kingdom feature compatible with a power-saving mode program. Moreover, transcriptional repression protects the genetic material of quiescent cells against intrinsic DNA damage that can occur as a result of transcription, especially if in the presence of agents capable of damaging DNA such as those that might occur during a transient exposure to noxious environmental agents [37, 38]. Despite the substantially decreased transcriptome size, we found that the levels of most transcripts in quiescent cells were closely correlated to those in proliferative cells (r up to 0.97), with most genes transcribed irrespective of the condition. This contrasts with yeast quiescent models in stationary phase cultures where up to 10% of all genes were transcribed in quiescent cells only and 27 % of all genes were exclusive to proliferative cells [39]. This may relate to the apparent absence of Pol II promoters in *Leishmania*.

Although the transcriptome size was diminished in all quiescent states measured compared to proliferative cells, the transcriptome composition revealed particular mRNAs whose proportion increased or decreased relative to others during quiescence. Since the transcriptome has a moderate correlation with levels of protein expression [40, 41], comparative transcriptome composition can provide insights into the biological processes associated with quiescence. We specifically sought changes found in quiescent cells irrespective of the life cycle stage or stimulus, and using GO/GSEA analysis identified three main functional categories as being upregulated: autophagy, proteolysis associated with leishmanolysin (GP63), and amastin/amastin-like proteins. An active autophagy process is a trait shared among quiescent eukaryotic cells that facilitates the non-specific lysosomal degradation of cytoplasmic components (microautophagy) and the recycling of unnecessary or damaged organelles, protein aggregates, lipid droplets, and lysosomes in a cargo-specific manner (macroautophagy) [42, 43]. We identified ATG8/AUT7/APG8/PAZ2 as the gene with the highest levels of TPM upregulation. ATG8 forms part of the LC3 conjugation system that is involved in the steps of pre-phagosome elongation, cargo recognition, and recruitment in macroautophagy [44]. Interestingly, one copy of the gene encoding an ‘FYVE zinc finger containing protein’ was also upregulated in all quiescent conditions. This protein forms part of the ATG12 conjugation system that facilitates the incorporation of the LC3 complex into the growing phagophore [45]. The enrichment of transcripts for ATG8 and FYVE suggests macroautophagy remains very active during quiescence. It was previously shown in *L. major* promastigotes that the number of cells containing autophagosomes, as well as the average number of autophagosomes per cell increases during the transition from logarithmic to stationary cultures. Moreover, mutants with impaired autophagy were defective in their ability to differentiate to non-proliferative metacyclic forms and less able to withstand starvation [46]. In yeast, it has also been shown that autophagy is increased during transition to a quiescent form [47]. During starvation, autophagy allows the recycling of crucial nutrients and under drug pressure or other environmental stress, the process can remove unnecessary organelles [48]. Indeed, *L. major* metacyclics recovered from the nutritionally challenging environment of stationary phase have a significantly reduced number of organelles compared to proliferative cells [49], pointing to an active mechanism of macroautophagy for their degradation.

Transcripts encoding GP63 were also upregulated across quiescent conditions. GP63 is a plasma membrane localized metalloprotease in both promastigotes and amastigotes [49], with secondary localisations within the reticulum endoplasmic, flagellar pocket, and with a secreted form known too [50, 51]. In promastigotes, GP63 increases during the transition to the stationary phase and is abundant in the infective non-proliferative metacyclics [52, 53]. Upon transmission to a mammalian host, GP63 plays a key role in the evasion and inactivation of the host innate immune system [54]. In *L. major*, GP63 modulates the normal course of phagocytosis by hampering the acidification of the phagolysosome [55], and it also prevents the ROS burst by proteolytically impairing the assembly of the NOX2 complex [54]. In amastigotes, GP63 is critical for their survival [51], being abundant in the flagellar pocket which is the major site of exocytosis [51]. Amastigotes secrete GP63 to cleave host proteins such as fibronectin in the extracellular matrix that in turn decreases the production of ROS by other parasite-infected macrophages [56]. Thus, GP63 upregulation in quiescent cells could be related to its many mechanisms to protect against innate immune responses to facilitate an effective infection and long-term survival within the mammalian and insect hosts [57, 58].

The other substantially upregulated gene set common to all quiescent forms corresponded to the amastins, a family of genes encoding cell membrane localised glycoproteins which are related to claudins that associate with tight junctions in metazoa [59]. Amastins were initially discovered in *Leishmania* amastigotes, however they are are also expressed in the insect form of trypanosomatids including the epimastigote of *Trypanosoma cruzi*, and are upregulated in metacyclic promastigotes of *L. infantun* isolated from the sand fly [60, 61]. Although the molecular function of amastins is unknown in *L. braziliensis* their knockdown decrease the infectivity of promastigotes towards macrophages *in vitro* and *in vivo* [59]. Amastin surface like-proteins (ALSP), low molecular weight proteins analogous to amastins, were also upregulated during quiescence. In *L. donovani* amastigotes ALSPs have been proposed to function as triglyceride lipases generating glycerol and fatty acids that can be transported into the parasite to support energy generation and the synthesis of specialised lipids [62]. Intriguingly, despite their name ALSP have a cytoplasmatic localization near to the flagellar pocket and, as is true for other lipases in *Leishmania*, they can be secreted. In *Saccharomyces*, quiescent cells under starvation accumulate triacyl glycerides within lipid droplets that can be metabolized to fatty acids by TAG lipases for the rapid resumption of growth following the replenishment of nutrients [63]. It will be of interest to see if a similar role of ALSP can be uncovered in *Leishmania*.

Metabolomics analysis also revealed quiescent cells had a decrease in the overall abundance of low molecular weight metabolites being among the most representative nucleotides and amino acids. These results are consistent with the GSEA that identified quiescent cells compared to those in proliferation downregulate transcripts involved in the metabolism of nucleotides and many amino acids *de novo* and salvage. Similar downregulation in amino acids was already reported in *L. brazileinsis* and *L. mexicana* quiescent amastigotes, a trait that matches their diminished protein synthesis and turnover [5, 19]. The downregulation in the levels of amino acids under PAT pressure is remarkable as they were still in an environment without nutrient limitations suggesting an active negative regulation instead of the lower levels resulting from a lack of carbon or nitrogen sources for their synthesis or scarcity for their salvage. A closer inspection in promastigotes and amastigotes revealed that at a transcriptional level only quiescent cells under PAT pressure downregulate four proteins related to the amino acid permease family, which suggests that decreased amino acid intake could be an important mechanism of regulation. A low level of amino acids is a hallmark of quiescence among bacteria and in *Mycobacterium*, the addition of glutamine sensitises quiescent forms to rifampicin since it alters their TCA cycle [64]. Against the overall down trend, quiescent cells across conditions had a clear increase in polyhexoses inferred to be part of the mannogen complex in these parasites, and a carbohydrate reserve already shown to be increased in stationary promastigotes and quiescent amastigotes of *L. mexicana* [32, 65]. The genome annotation of genes involved in the metabolism of mannose is not yet complete in *Leishmania* models. Very recently *L. mexicana* knockout lines for an array of multicopy genes encoding 7 ‘mannosyltransferase/phosphorylases’ (MSPs) have proved them essential for the survival of promastigotes under stress induced by elevated temperatures or acidification of the medium as well as for the survival of amastigotes *in vivo* [65]. Although MSPs are not yet annotated in the reference genome used in this study, we identified several modulated transcripts involved in the metabolism of mannose in a condition-specific manner (Table S2f). The gene GDP mannose 4-6 dehydratase (GMD) is of particular relevance because it was significantly upregulated in all quiescent cells across conditions (although only more than 2 fold in stationary promastigotes and amastigotes). GMD participates in the conversion of GDP-mannose to GDP-fucose, a metabolite that has been found to be essential for the survival of *L. major* [66]. As GDP-fucose is a source of sugar for glycosylation, the availability of mannose may be relevant not only as an alternative source of energy but also as a donor for the synthesis of glycosylated proteins. In *Trypanosoma brucei* a conditional null mutant for GMD has shown GMD is essential for the long term survival of both the procyclic and bloodstream forms [67]. Quiescent cells also had a trend of increased levels of free fatty acids despite the downregulation of transcripts associated with the fatty acid biosynthesis across all quiescent conditions. This is not all surprising as *Leishmania* can switch from *de novo* synthesis to salvage from the host (or the medium *in vitro*) for the maintenance of their pool of fatty acids and lipids [68]. Moreover, the potential of a link between ALSP and the levels of fatty acids is another possible association between our transcriptomics and metabolomics findings that requires further exploration.

Our study shows common traits of *Leishmania* quiescent cells that contribute to the understanding of the maintenance of quiescence at molecular and metabolic levels regardless of the stimuli inducing this phenotype. We show for the first time quiescence can be induced in response to antimonial drug pressure. This will have a big impact in the field of drug development as new therapies should prove to be effective against the resilient quiescent forms. Considering the limited resources for drug development we recommend that drug targets with evidence of being essential for proliferative and quiescent forms should be ranked first. Combination therapies between existing drugs and a metabolite causing a metabolic unbalance in quiescent subpopulations like in the case of glutamine plus rifampicin for *Mycobacterium* could be also tested for *Leishmania*. It will also be of interest to investigate if the capability to become quiescent in response to drug pressure varies among clinical isolates and if quiescent features could be associated with therapeutic failure.

## 4. Materials and Methods

### 4.1. Leishmania clinical isolate and cell culture

The clinical isolate MHOM/PE/04/PER091 (provided by the Institute of Tropical Medicine Alexander von Humboldt) was used to generate the transgenic clone MHOM/PE/04/PER091 EGFP Cl1 on which all the further experimental work was performed. Promastigotes were maintained in complete M199 (M199 medium at pH 7.2 supplemented with 20% fetal bovine serum, hemin 5 mg/L, 50 μg/mL of hygromycin Gold, 100 units/mL of penicillin and 100 μg/mL of streptomycin) at 26 °C with passages done twice per week. To obtain axenic amastigotes, 1 mL of stationary promastigotes was centrifuged (1500 g, 5 min) and the pellet was re-suspended in 5 mL of complete MAA (M199 at pH 5.5, supplemented with 20% fetal bovine serum, glucose 2.5 g/L, 5 g of tryptic soy broth, hemin 5 mg/L, 25 μg/mL of hygromycin Gold, 100 units/mL of penicillin and 100 μg/mL of streptomycin) and incubated at 34 °C. Morphological amastigogenesis was observed at day 3 and the strain was maintained indefinitely as amastigotes with passages done twice per week. Samples for experimental analysis were prepared in complete M199 and MAA but without hygromycin. For the preparation of parasites under drug pressure, an equal volume of exponentially growing parasites were mixed with complete M199 or MAA containing 2 μg/mL of potassium antimonyl tartrate trihydrate (PAT) (sigma Aldrich 383376). The final concentration represented ~10 × the PAT IC50 in promastigotes as measured by the resazurin test.

### 4.2. Development of an EGFP clonal line

Enhanced GFP (EGFP) was integrated within the 18S ribosomal DNA locus with the use of the pLEXSY-neo2 system (Jena Bioscience) as previously reported elsewhere [6, 69, 70]. A clonal line was obtained from the transgenic rEGFP parasites with the ‘micro-drop’ method [6].

### 4.3. Promastigotes and amastigotes enrichment with Ficoll and Percoll gradients

*Leishmania* cells with good viability were enriched with Ficoll and Percoll gradients [71]. Briefly, promastigotes in stationary phase or after 48 hrs of PAT pressure were harvested and centrifuged at 2000 g for 5 min. The pellet was resuspended in 4 mL of non-supplemented M199. The suspension of cells were added as the last layer of a Ficoll gradient (Ficoll Type 400) prepared in 15 mL tubes containing 2 mL of Ficoll 20% at the bottom and Ficoll 10 % on top. Gradients were centrifuged at 1300 g for 15 min at RT. After centrifugation, the fraction of Ficoll 10% was recovered and washed with 10 mL of cold PBS. After centrifugation at 2500 g for 5 min the pellet was resuspended in 1 mL of cold PBS. Cell numbers per mL were calculated and aliquots for both RNAseq (2 × 10 ^7^cells) or metabolomics (4 × 10 ^7^cells) were prepared. The pellets for RNAseq were stored at −80 °C while samples for metabolomics were processed immediately after harvesting. An extra aliquot of each sample was taken for monitoring of EGFP expression and cell viability. Amastigotes in stationary phase or after 48 hrs of PAT pressure were harvested and centrifuged at 2000 g for 5 min. Viable amastigotes in stationary phase or under drug pressure were enriched with a Percoll gradient. Briefly, cells were harvested and resuspended in 5 mL of M199. The cellular clusters were disrupted by passing the medium through a 26 G needle over 3 times. The suspension was centrifuged at 2000 g for 5 min and the pellet was eluted in 6 ml of Percoll 45 %. The gradient was formed by placing 1.5 mL of Percoll 70 % at the bottom of a 15 mL tube, overlayed with the suspension of cells in Percol 45 % and a final gradient of Percoll 25% on top. The gradient was centrifuged at 2300 g for 45 min at 4 °C. The fraction of Percoll 45 % and the intersection with Percoll 70 % was recovered and washed with 10 mL of cold PBS. After centrifugation, the pellet was resuspended in 1 mL of cold PBS. Counting and aliquots were prepare as in promastigotes.

### 4.4. Monitoring rEGFP expression and cell viability by flow cytometry

rEGFP expression and cell viability at single cell level were monitored by flow cytometry. Briefly, EGFP expression was quantified as an indicator of quiescence while the cell viability was evaluated by using the NucRed dead 647 (Thermo fisher scientific) for the staining of dead cells and Vybrant Dye Cycle violet (Thermo fisher scientific) for the staining of cells with nucleus and kinetoplast. The samples were analysed with a calibrated flow cytometer BD FACS VerseTM in the medium flow rate mode. A wild-type (non rEGFP) line and a non-stained sample were included in each experiment as negative controls. In order to compare the relative fluorescence units (RFU) among samples the acquisitions were made with the same settings during all the experiments. The FCS files were analyzed with the FCS 5 express Plus research edition. Gates were created as described elsewhere [6], with an additional step where a gate for cells negative for NucRed but positive for Vybrant Dye cycle violet (Vybrant DC) was created. The levels of rEGFP expression were evaluated in the populations of viable cells being NucRed ^-^ and Vybrant DC ^+^.

### 4.5. RNA and library preparation

An amount of 2 × 10 ^7^ parasites per condition and stage were harvested, and the total RNA was isolated with the RNeasy Micro Kit (Qiagen). The RNA was eluted in a total volume of 22 μL and quantified with the Qubit^™^ RNA BR Assay Kit. The integrity of the RNA was evaluated by electrophoretic separation with the RNA ScreenTape system (Agilent). For absolute normalization (to correct for the differences in total RNA content per cell across conditions), samples were divided into two groups and mixed with 1 μl (pre eluted 1:500 in H20) of one of two predefined mixtures of external RNA controls for which their identity and fold change when comparing Mix1/Mix2 are known (ERCC ExFold RNA Spike-In Mixes). Strand-Specific RNA-Seq service was provided by Genewiz where up to 250 ng of RNA was used to prepare the library with a polyA selection and the NEB-next ultra II directional RNA library preparation kit. The library was then sequenced using Illumina NovaSeq 2 × 150 pb sequencing configuration.

### 4.6. RNAseq data analysis

Fastq files were analysed using Chipster server. Briefly, The reads were mapped to a combined genome containing the reference nuclear and kinetoplast (maxicircle) genome of *L. braziliensis* (MHOM/BR/75/M2904) reported by Gonzales et al 2018 [72] and the ERCC spikes-in using the BWA aligner *mem* command with default parameters. The aligned reads were counted with HTSeq using the htseq-count command considering the features ID from all the CDS in the kinetoplast maxicircle (18), nuclear genome (8277), and ERCC spikes-in (92). After evaluating the transcriptome coverage, that ranged between ~19 X in Ama Log_PAT to ~65 X in Ama Log (Supplementary Table S1b), the raw counts table was processed with two approaches.

Firstly, relative quantification representing differences in the number of molecules of each transcript per cell were calculated. For this purpose, the raw counts table was normalized using the counts coming from the external ERCC spikes-in controls. This step was done independently for all samples having the same ERCC Spikes-in mixture (Mix 1 or Mix2). Briefly, for each spike a positive detection across samples and a cutoff of minimum 5 reads per sample were used as parameters to filter out spikes with low and likely random number of reads. Subsequently, the per-sample normalization factor was obtained with the following formula:

NFz= ∑(read count of spike i in sample Z / median read count of spike i over all samples) /n. Where n represents the number of spikes passing the prefiltering.

The normalized count were then analysed with the DESeq2 package. The size factors were calculated using the ERCC spikes-in belonging to the group B (spikes with expected log2 FC equal to 0 across samples) and the formula for the negative binomial model included the condition and stage of the parasite as explanatory variables including also an interaction term for both factors. Differences between dichotomic comparisons were considered significant when Log 2 FC was higher than 1 or lower than −1 with a Benjamini-Hochberg adjusted *p* value lower than 0.05. The VennDiagram package was used to generated plots that show the interrelationship of modulated genes across conditions.

Secondly, we evaluated the transcriptome composition by performing relative quantification representing the abundance of each transcript with respect to its own transcriptome. For this purpose, the data was normalized with the TPM method (transcripts per million). The following formula was used: TPM= (reads mapped to transcript/ transcript length) × 10^6^/ ∑ (reads mapped to transcript/ transcript length). The data was then analysed with the DESeq2 package. Because the normalization was already done, the normalization factor we set to 1 during DEseq2 calculations. The formula and criteria for significant differences were as described above.

### 4.7. Transcriptomics functional enrichment and network analysis

Gene ontology (GO) and Gene Set enrichment analysis were performed to identify which functional categories were modulated in quiescent conditions compared to proliferative cells. g:Profiler tools were used to map the list of genes with modulated TPM to known gene ontology terms [73]. g:Profiler performs a cumulative hypergeometric test and multiple testing correction (Benjamini-Hochberg FDR) to detect statistically significantly enriched biological processes, pathways, regulatory motifs, and protein complexes. The statistical domain scope was set to all known genes and the input data consisted of independent lists of genes with either upregulated TPM (adjusted *p* < 0.05, log2 FC > 1) or down regulated TPM (adjusted *p* < 0.05, log2 FC < −1). The summarization and network analysis for the significant GO terms with an adjusted *p* < 0.05 were performed with Revigo and Cytoscape [74, 75], respectively. The threshold for redundancy reduction was set to small (0.5). Per each condition, GSEA was performed using GAGE version 2.42.0 and the list of log2 FC of all genes [76]. Two independent One sample Z tests were performed, one for upregulation and another for downregulation. A gene set was considered significantly modulated if the global *p* value for the One sample Z test was lower than 0.05. Data for individual tests across conditions were then integrated as follow: up or down in all if the *p* values for a particular set across conditions were <0.05. Unchanged in all if the *p* values across conditions were > 0.05 and other for sets without significant change across conditions. After integration, the changes for each set were visualized in a heatmap format.

### 4.8. Metabolomics

Per each condition, 4 biological replicates were prepared and an amount of 4 × 10^7^ cells were harvested for metabolites extraction with a mixture of CHCl_3_/MeOH/H_2_O (1:3:1) as previously described (Berg et al., 2015). Samples were analysed with Liquid chromatography and mass spectrometry using an Orbitrap Exactive mass spectrometer (Thermo Fisher) coupled to a 2.1mm ZIC-HILIC column at Glasgow Polyomics (University of Glasgow, Scotland). Two quality control samples were included: 1) Authentic standard mixes containing in total 178 metabolites (76–516 Da) representing a wide array of metabolic classes and pathways and 2) pooled sample of all extracts for which MS2 fragment analysis was also performed. The peaks were detected and annotated with IDEOM [77]. Subsequently, features were visually verified and filtered out when they had multiple peaks or a not well-defined shape. The feature with the highest average intensity across samples was kept when a putative metabolite had duplicated entries. The TIC was calculated per sample as the sum of all annotated features. The clean data was analysed with two methods of normalization. The first, used as reference the input of cells that was equivalent in all conditions. Therefore, further process was not needed. The second method adjusted the intensities of each feature by the TIC of its corresponding sample. For the purpose of differentiation, we will refer to this method as IPT. After normalization, all missing values were replaced by 1/5 of the minimum value across samples and transformed to a logarithmic scale. The metabolomics package was used to perform the statistical analysis and calculate the fold changes [78]. Briefly, within each stage and for each metabolite, the mean of each quiescent condition was compared to the mean of the proliferative condition using a paired T-Test. The Benjamini-Hochberg method was used to control for the false discovery rate and calculate the adjusted *p* values. Metabolites changes were considered to be significant if they had a |log2 FC| >1 and a BH adjusted *p* value <0.05.

## Supporting information

JaraM_Leish_quies_supplementary_figure

JaraM_Leish_quies_Supplementary_tables

## Supplementary Materials

The following are available online, Figure S1: Conditions Sta and Log_PAT drop their total RNA content, Figure S2: Experimental outline for the transcriptomic analysis, Figure S3: Normalization of samples with the use of ERCC spikes-in, Figure S4: Overview of the 135 genes with modulated TPM in all quiescent conditions, Figure S4: Quiescent cells have changes in the composition of their metabolome when compared to proliferating cells, Table S1: Transcriptome ERCC spike in normalized, Table S2; Transcriptome TPM normalized, Table S3: Metabolomics with Cell and IPT normalization.

## Author Contributions

Conceptualization, M.A.D., J-C.D., M.J.; methodology, M.J, I.M.; software, C.R., H.I., M.J.; validation, M.J.; formal analysis, C.R., H.I., M.J.; investigation, M.J., I.M.; resources, J-C.D., M.B., M.A.D., M.J.; data curation, C.R., M.J.; data interpretation M.J., M.B.; writing—original draft preparation, M.J.; writing—review and editing, M.B., J-C.D., M.A.D.; visualization, M.J.; project administration, M.J.; funding acquisition, M.A.D., J-C.D., M.J. All authors have read and agreed to the published version of the manuscript.

## Data Availability Statement

Raw RNAseq data is available in the Sequence Read Archive under project accession number PRJNA767428 (https://dataview.ncbi.nlm.nih.gov/object/PRJNA767428?re-viewer=3ls9fdjd87dn2d70ss61gi0hk2)

## Acknowledgments

We acknowledge the LeishNat-Drug-R [contract ICA4-CT-2001-10076] and professors Alejandro Llanos and Jorge Arevalo from the Institute of Tropical Medicine Alexander von Humboldt for providing us the clinical isolate used to derive that transgenic clonal line PER091 EGFP Cl1. MPB was funded as part of the Wellcome Trust core grant to the Wellcome Centre for Integrative Parasitology (grant 104111/Z/14/Z). Acknowledgements to Pieter Monsieurs for the edition and proofreading of the section of materials and methods.

