## Supplementary material for "Transcriptional shift and metabolic adaptations during *Leishmania* quiescence using stationary-phase and drug pressure as models": JaraM_Leish_quies_supplementary_figure

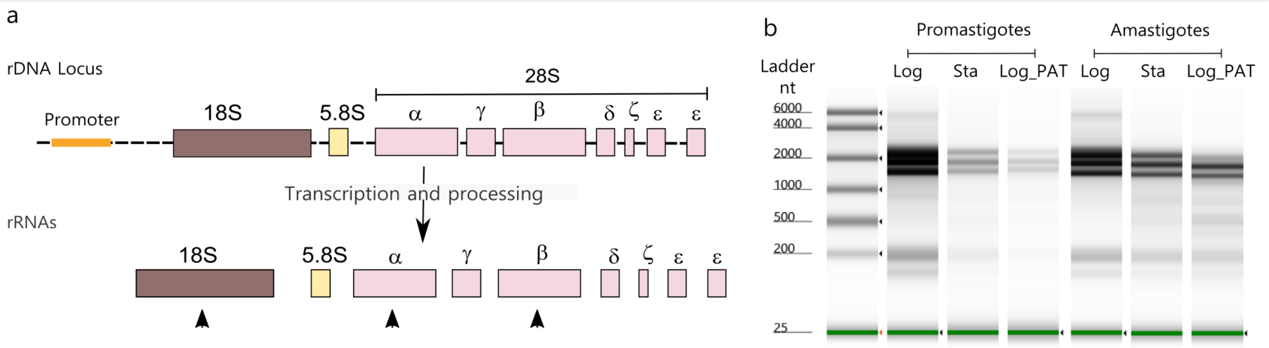


**Figure S1.** Conditions Sta and Log_PAT drop their total RNA content. a, Outline of the unique rRNAs processing in Trypanosomatids. The 28S rRNA that is an unique fragment in most of the eukaryotes is fragmented into 7 individual mature rRNAs. Because this difference in the processing, three main bands (corresponding to the rRNAs with arrows) instead of most common pattern of two, can be visualized in a electrophoretic run of total RNA. b, Electrophoretic analysis of the total RNA content showing the 3 expected bands for the rRNAs 18s , 28S β and 28S α. Each track was loaded with the total RNA corresponding to the same amount of cells (3.9 x 105). The image shows the substantial decrease in the total RNA content in conditions Sta and Log_PAT compared to proliferative cells and also the higher proportion of rRNAs over other RNAs across conditions.

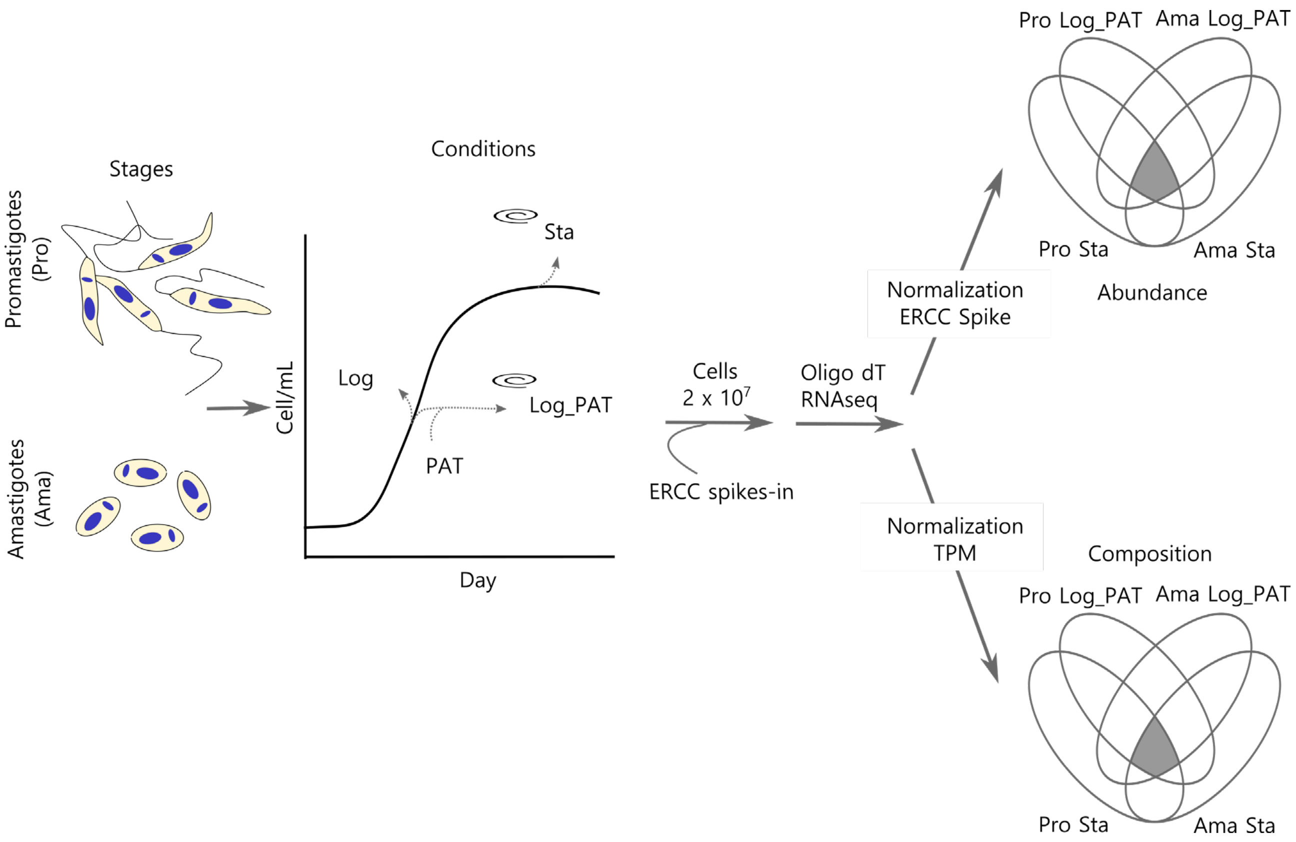


**Figure S2**. Experimental outline for the transcriptomic analysis. Promastigotes and amastigotes harvested during the exponential phase of the growth curve were used as a control for cells in a proliferative condition (Log). Quiescent cells were induced by stationary phase (condition STA) or PAT drug pressure (condition Log_PAT). Enrichment for cells with good viability through density gradients was performed for conditions Log and Log.PAT. A total of 2 x 107 cells in each condition were harvested and their total RNA was mixed with ERCC spikes-in as external RNA controls. RNAseq library preparation was done with an Oligo dT primer to target all mRNA. After Illumina sequencing, the raw reads were normalized with the ERCC spikes-in or the TPM method to evaluate the transcriptome abundance/cell and the transcriptome composition, respectively. Condition Log within each stage was always used as a reference to calculate the differentially expressed genes. The data of each dichotomic comparison was integrated to identify transcripts that were modulated and shared among all quiescent conditions ( up or down) regardless the stage or stimuli inducing quiescence (in the figure represented by all transcripts within the shaded area of the Venn Diagram). PAT; potassium antimonyl tartrate.

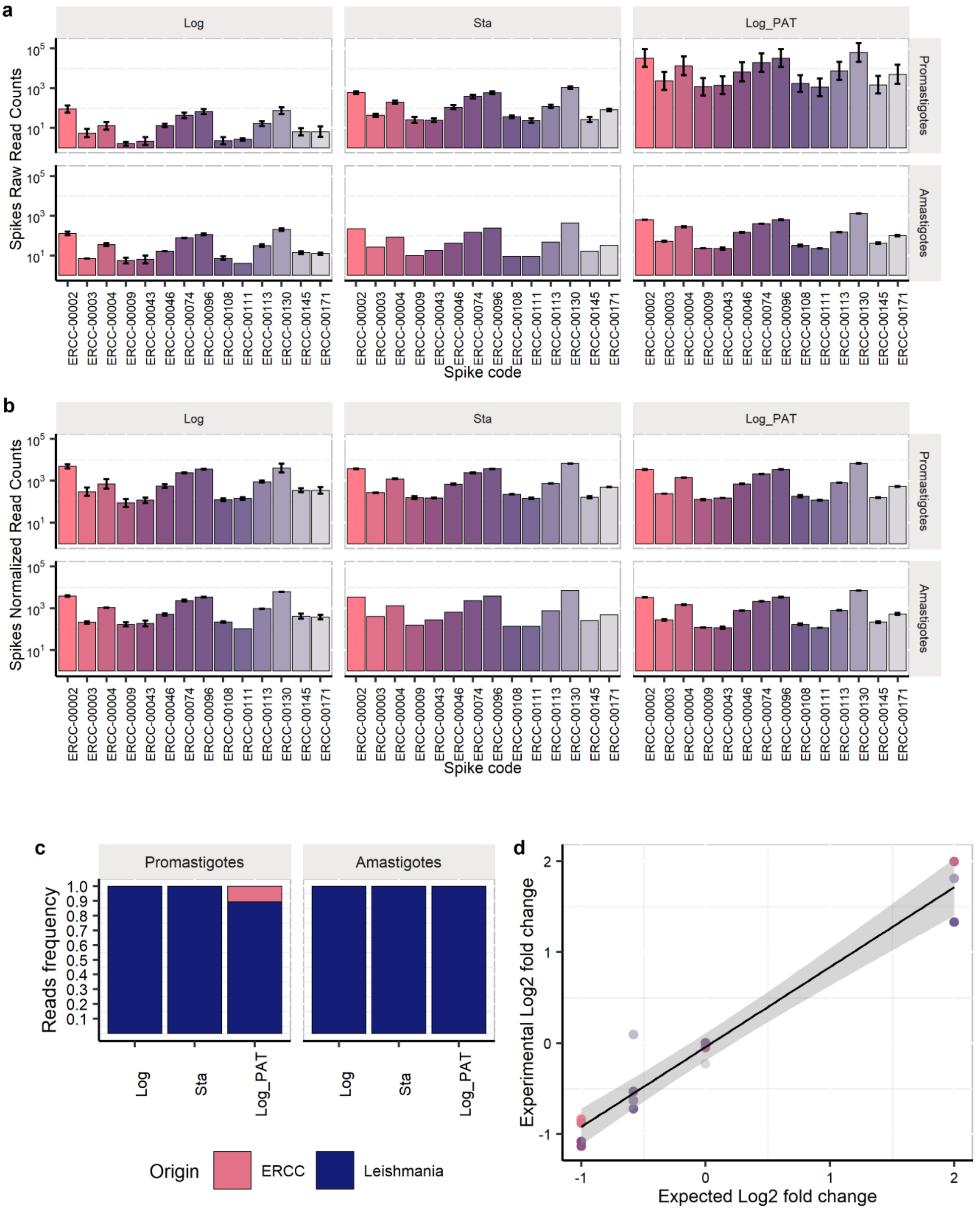


**Figure S3.** Normalization of samples with the use of ERCC spikes-in. A) Raw read counts for the ERCC spikes-in before normalization. An equal amount of ERCC spikes-in per each 2 x $10^7$ cells was added to each sample. Despite each sample having the same amount of ERCC spikes-in, the raw read counts for individual ERCC spikes-in differed among samples. These differences evinced the bias introduced by the differences in RNA content/cell, the library preparation and sequencing. B) Normalized read counts for the ERCC spikes-in. After the per sample normalization factor was calculated and applied to each sample, the resulting normalized read counts for each ERCC spikes-in had very similar values across samples. This process allowed us to quantify differences in the abundance of mRNAs per cell among the different conditions. In samples with the same spikes mixture, only ERCC spikes-in detected in all samples with raw read counts >5 were considered to calculate the normalization factor. C) Frequencies of reads coming from the ERCC spikes-in and *Leishmania*. D) There was a good correlation (0.97) between the experimental log2 fold change and the expected log2 fold change for the ERCC spikes-in used for the normalization. The experimental Log2 FC was calculated by dividing the mean of normalized read counts for each spike in samples seeded with the mix1 and the mean of normalized read counts of each spike in samples with the mix2.

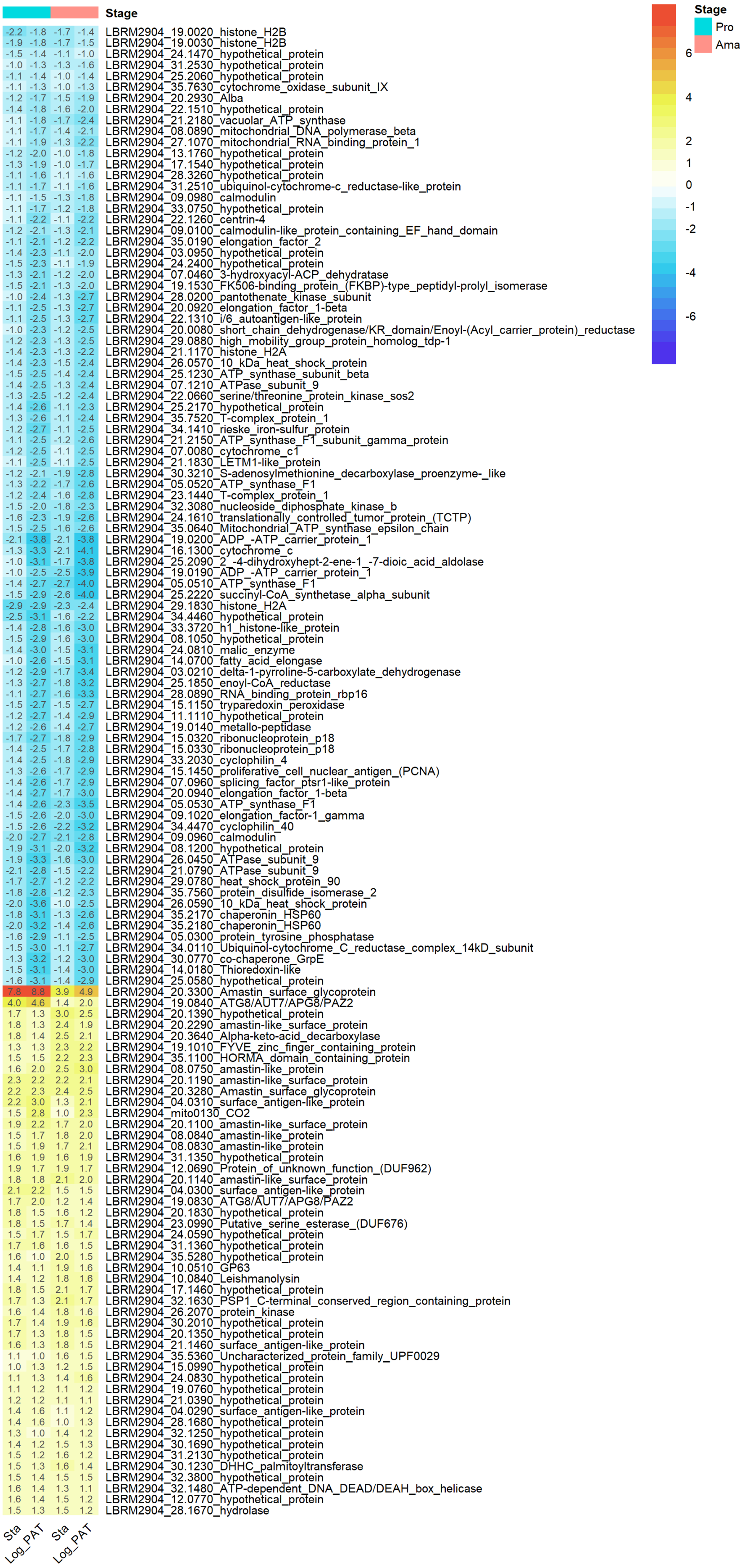


48

49 **Figure S4**. Overview of the 135 genes with modulated TPM in all quiescent conditions. The numbers
50 and color key represents the Log2 FC in quiescent cells compared to proliferative cells.

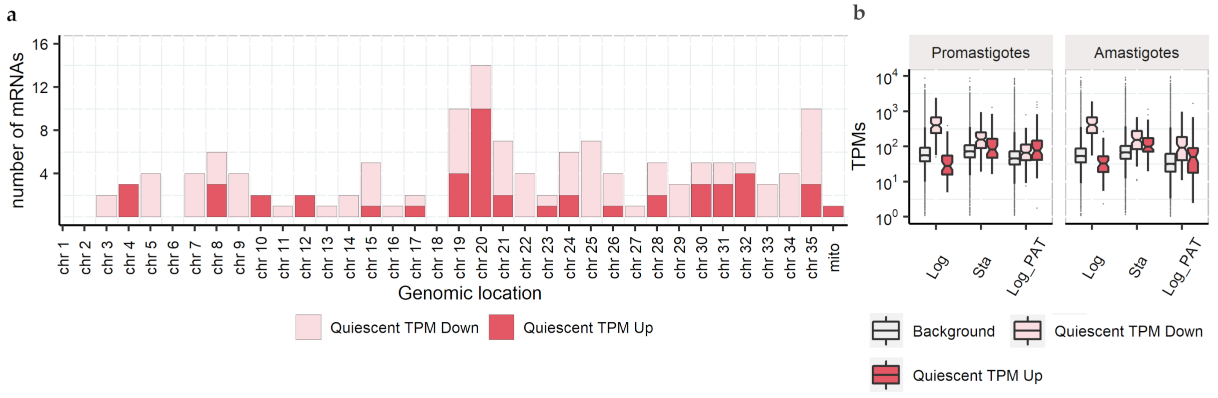


**Figure S5.** Transcripts commonly modulated across quiescent condition are located in many genomic locations and their TPMs distribution differs of the overall transcriptome. a, Genomic location for the core set of 135 mRNA with modulated TPMs across all quiescent conditions. The bars show the number of modulated mRNAs mapping to a specific chromosome (chr) or the mitochondrial maxicircle (mito). b, Box plots showing the broad distribution of TPMs for individual mRNA (TPMs, y-axes) across conditions. In each *Leishmania* condition, the pool of mRNAs consists of a broad range of transcripts, with some having many copies (high TPM values) and others with few copies (low TPM values). In proliferative promastigotes and amastigotes the TPMs distribution for the core sub set of mRNAs modulated in all quiescent cells (Quiescent TPM Down=87, Quiescent TPM up= 48) differs of the distribution for the overall pool of mRNAs (Background). Quiescent TPM Down corresponds to transcripts that initially have many copies (or highly repetitive) in proliferative cells, as their distribution is above the median of the overall pool of mRNAs. The opposite is true for the set Quiescent TMP up, as they initially had low copy number in proliferative cells. Each box represents the 1st quartile (25th percentile) and 3rd quartile (75th percentile), the horizontal line represents the median, and both whiskers represent 1.58 times the Interquartile range.

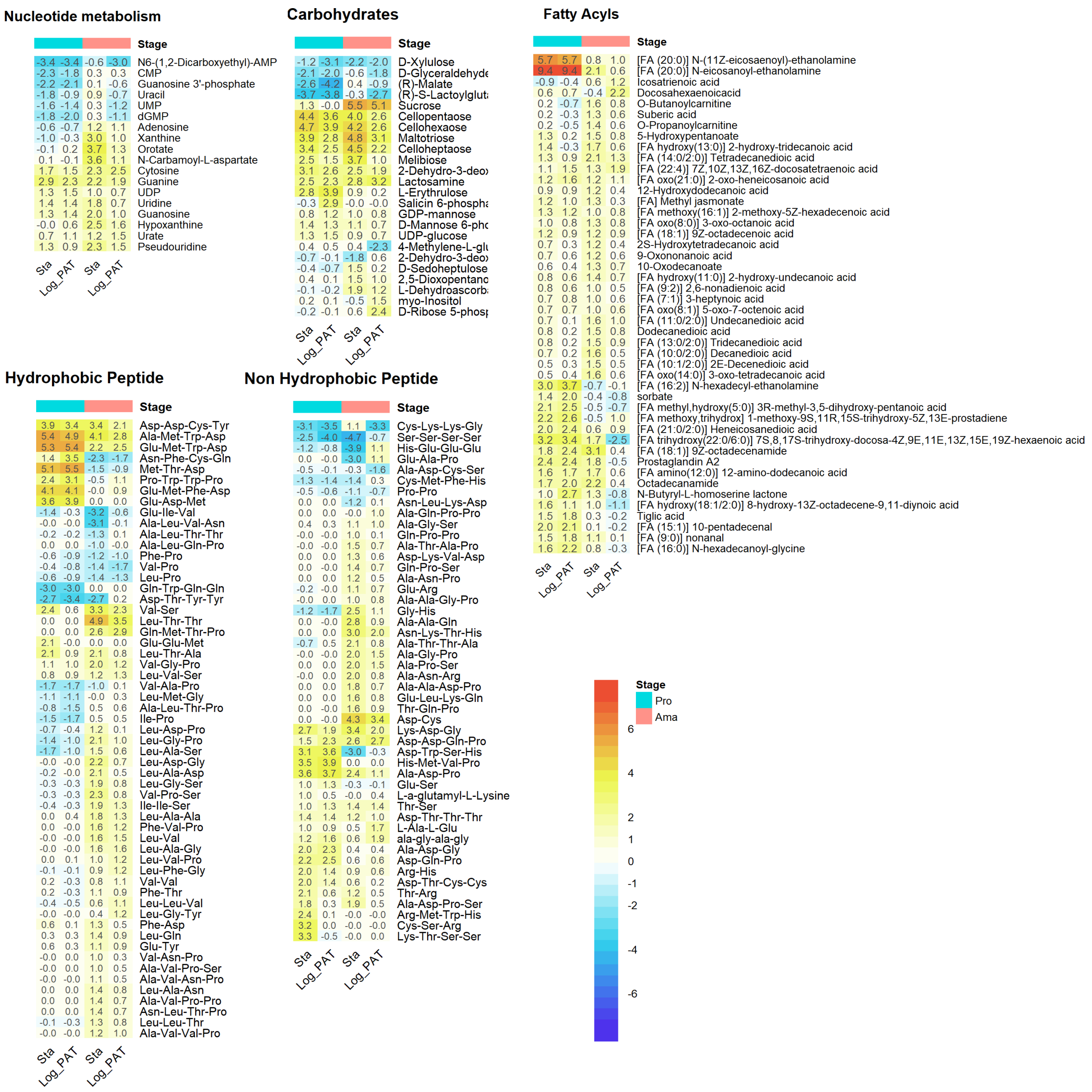


**Figure S6.** Quiescent cells have changes in the composition of their metabolome when compared to proliferating cells. The fatty acyls were the category showing a clear trend of overall upregulated levels. The numbers represent the Log2 FC (IPT normalized dataset). On each category, only metabolites for which at least one of the conditions had | Log2 Fold change| > 1 and a BH adjusted P <0.05 are shown. Pro, promastigotes; Ama, amastigotes.
